# Engineering and characterization of a long half-life relaxin receptor RXFP1 agonist

**DOI:** 10.1101/2022.04.19.488796

**Authors:** Sarah C. Erlandson, Jialu Wang, Haoran Jiang, Howard A. Rockman, Andrew C. Kruse

## Abstract

Relaxin-2 is a peptide hormone with important roles in human cardiovascular and reproductive biology. Its ability to activate cellular responses such as vasodilation, angiogenesis, and anti-inflammatory and anti-fibrotic effects have led to significant interest in using relaxin-2 as a therapeutic for heart failure and several fibrotic conditions. However, recombinant relaxin-2 has a very short serum half-life, limiting its clinical applications. Here we present protein engineering efforts targeting the relaxin-2 hormone in order to increase its serum half-life, while maintaining its ability to activate the G protein-coupled receptor RXFP1. To achieve this, we optimized a fusion between relaxin-2 and an antibody Fc fragment, generating a version of the hormone with a circulating half-life of up to five days in mice while retaining potent agonist activity at the RXFP1 receptor both *in vitro* and *in vivo*.

## Introduction

Relaxins are small protein hormones belonging to the insulin superfamily, exerting a variety of biological activities through the activation of G protein-coupled receptors (1). Within this family is relaxin-2, a reproductive hormone responsible for mediating many of the physiological changes of pregnancy through its cognate receptor, RXPF1 (2). Relaxin-2 signaling through RXFP1 leads to vasodilation, angiogenesis, collagen degradation, and anti-inflammatory effects. In addition to relaxin-2’s role in pregnancy, these cellular responses also regulate the physiology of multiple organs in both sexes, including the liver, kidney, heart, lungs, and blood vessels (3,4). Activation of the pleiotropic effects downstream of RXFP1 can improve cardiac function and decrease fibrosis levels, which has generated interest in using relaxin-2 as a treatment for cardiovascular and fibrotic diseases (5–7).

In animal models, recombinant relaxin-2 (serelaxin) has yielded promising results for the treatment of heart failure and fibrosis of the liver, lungs, kidneys, and joints (8–13). However, in large-scale clinical trials for acute heart failure, serelaxin treatment did not significantly decrease patient rehospitalization or mortality, although patients showed some short-term relief of symptoms such as dyspnea (14). One potential cause of these results is the short serelaxin administration time of 48 hours, while patient data was collected up to 180 days after treatment. As a small protein hormone, serelaxin is rapidly cleared from circulation, with a serum half-life of less than 5 hours (15). Therefore, the beneficial effects may be lost relatively quickly without continuous or repeated intravenous administration, limiting the use of serelaxin in many chronic conditions and introducing challenges around patient compliance.

The native relaxin-2 molecule has a two-polypeptide chain structure, with an A-chain and B-chain connected by disulfide bonds, structurally similar to insulin (16). Protein engineering of the relaxin-2 molecule and small molecule screening have each been explored to develop agonists of RXFP1 beyond the highly potent native relaxin-2 peptide, which has an EC_50_ of around 100 pM (**Figure 1c, Table S1**). Small molecule screens have proven to be challenging (17), and only one series of small molecule agonists have been reported, with an EC_50_ of approximately 100 nM for the lead molecule, ML290 (18,19). Versions of relaxin’s B-chain have been produced and tested for activity at RXFP1, however, the B-chain alone shows low signaling potency, with an EC_50_ of around 7 μM (20). Optimization of B-chain only variants and the addition of lipid modifications resulted in peptides able to activate RXFP1 with improved potency and half-lives of up to 9 hours (21). Additionally, studies with mouse models of heart failure have recently tested a fusion between relaxin-2 and an antibody Fc, however no details of the protein sequence or engineering methods were reported (22).

**Figure 1:**
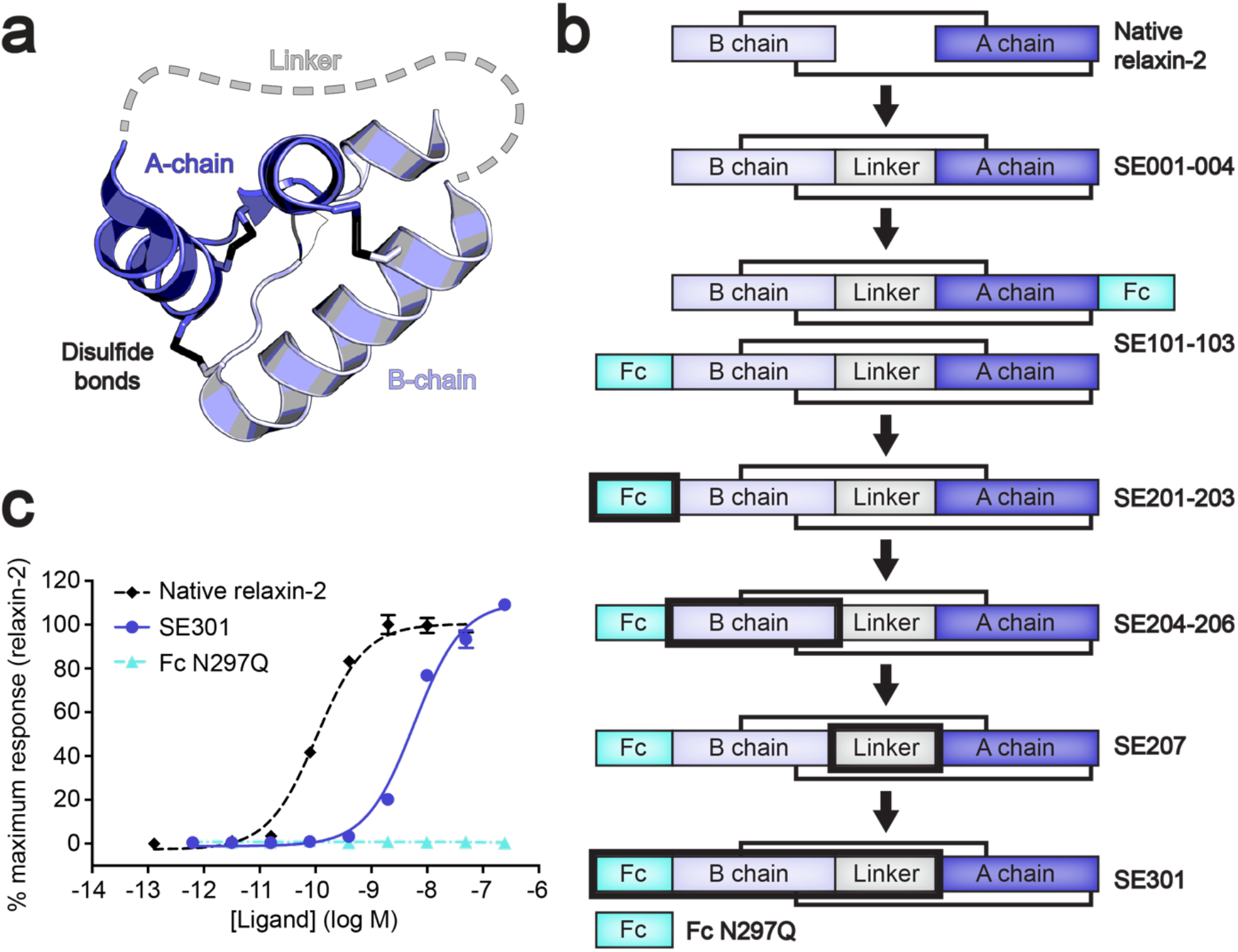
Engineering of an Fc–single-chain relaxin-2 fusion. **a**, Diagram of single-chain relaxin-2 using the X-ray crystal structure of human relaxin-2 (PDB ID: 6RLX) (16). **b**, Overview of the rounds of optimization for the Fc–relaxin-2 fusion. The black box highlights regions that were optimized in each iteration. **c**, CRE-SEAP Gs signaling assay data for human RXFP1 using SE301 and Fc N297Q compared to native relaxin-2. Data are normalized to the native relaxin-2 response and are mean ± s.e.m. from technical triplicates.

Creating a fusion with an antibody Fc fragment is an established method of increasing the serum half-life of a protein of interest. Through the neonatal Fc receptor (FcRN) binding, antibodies can avoid lysosomal degradation and be recycled back into circulation, resulting in a half-life of two to three weeks for most IgGs (23). Fc fusions to a protein of interest allow them to take advantage of the same recycling mechanism, conferring a long serum half-life to a protein that would otherwise be cleared quickly from circulation (24,25). Given the two-chain structure of the relaxin-2 hormone, a simple fusion to an Fc fragment is impossible. Moreover, any modifications to the relaxin-2 protein must be weighed against reducing signaling potency. In order to engineer an Fc–relaxin-2 fusion protein, we created a single-chain relaxin-2 molecule using a linker to connect the two peptide chains. Optimizations of the single-chain relaxin-2 sequence and the fusion to human IgG1 Fc generated a molecule with high biochemical stability and yield. The optimized fusion, SE301, maintains a high level of biological activity while gaining a long serum half-life. Moreover, it is straightforward and cost-effective to produce in large quantities. Here we describe the rational design of the SE301 molecule and the characterization of its *in vitro* and *in vivo* activity and pharmacokinetics profile.

### Engineering of a single-chain relaxin-2

The native relaxin-2 hormone is translated as a single polypeptide chain. In order of sequence, the prohormone consists of a B-chain of 29 residues, a connecting C-chain of 108 residues, and an A-chain of 24 residues. After translation, the C-chain of prorelaxin-2 is cleaved out by proteolytic digestion. The resulting protein is the mature form of native relaxin-2, in which the B-chain and A-chain are separate alpha helical peptides connected by two interchain disulfide bonds (26).

The first methods of producing native relaxin-2 utilized chemical synthesis of the two separate chains in reduced forms, followed by an oxidative step to form the disulfide bonds (27,28). Methods using recombinant DNA technology were able to improve the yields for relaxin-2 using a construct containing a “mini-C” peptide linker between the B and A chains. These methods utilized the published X-ray crystal structure of relaxin-2 to design a shortened C-chain length of 13 residues that would still be long enough to connect the two chains (16,29). These relaxin proteins were expressed in *Escherichia coli,* purified from inclusion bodies, and refolded. The “mini-C” linkers were then removed by protease digestion, resulting in a recombinant form of the native relaxin-2 hormone (29).

To generate a mature single-chain version of relaxin-2, we first tested a linker originally used as a cleavable “mini-C” peptide for relaxin-3, another member of the relaxin hormone family (30). We expressed and purified the single-chain relaxin-2 with an N-terminal hemagglutinin signal sequence as a His-tagged secreted protein in mammalian cells without removing the “mini-C” linker (**Figure 1a,b**). The expression protocol was purposefully chosen to determine if a less labor-intensive method could be used to produce properly folded and biologically active relaxin-2. The protein showed a monodisperse size exclusion profile and high purity by Coomassie-stained SDS-PAGE gels (**Figure S1a,b**). To determine whether our single-chain relaxin-2 maintained activity at RXFP1, the protein was tested using a cell-based assay for Gs signaling. The single-chain relaxin-2 (SE001) maintained sub-nM potency at human RXFP1, illustrating the feasibility of generating single-polypeptide relxin-2 molecules with high biological activity from mammalian expression systems (**Figure S1c**).

In later rounds of protein engineering, the sequence of the “mini-C” linker was further optimized (**Figure 1b**). The first linker, Asp-Ala-Ala-Ser-Ser-His-Ser-His-Ser-Ser-Ala-Arg, contained several Ser residues and His residues that were removed in the redesigned sequence. Ser residues were changed to remove potential sites of O-linked glycosylation and His residues to prevent any pH-dependent changes in binding affinity. After redesign, the new linker consisted of the sequence Asp-Ala-Ala-Gly-Ala-Asn-Ala-Asn-Ala-Gly-Ala-Arg (SE207).

### Optimization of Fc–single-chain relaxin-2 fusions

The biologically active single-chain relaxin-2 created an opportunity to alter several of the protein’s properties through engineering additional fusions. The main purpose of these modifications was to lengthen the short serum half-life of the small relaxin-2 protein, in order to generate a long half-life agonist of the RXFP1 receptor. To accomplish this, we tested fusions of a human IgG1 Fc antibody fragment to either the N- or C-terminus of single-chain relaxin-2 (SE101-103, **Figure 1b**). G_s_ signaling assays showed that N-terminal fusions maintained higher signaling potency at human RXFP1 than C-terminal fusions (8.3 nM vs. 49.4 nM, **Figure S2a**). These proteins also had increased purification yields, up to 400-fold higher than the initial construct SE001.

At this stage, a mutation was introduced to the human IgG1 Fc fragment for all further constructs. The mutation, N297Q, removes a glycosylation site from the Fc fragment important for IgG1 effector functions. As a result, the abilities of IgG1 Fc to activate complement and antibody-dependent cellular cytotoxicity are ablated in the N297Q mutant (31). The fusion site between Fc N297Q and the N-terminus of single-chain relaxin-2 was the next region to be optimized (**Figure 1b**). The Fc fragment had initially been fused to the single-chain relaxin-2 N-terminus by a linker of Gly-Gly-Ser repeats. Fusions with a shorter 3-residue linker length achieved higher signaling potency than a 12-residue Gly-Gly-Ser linker (SE101-103, **Figure S2a**). Sequences for the 3-residue linker were then varied by constructing linkers with the sequence Ala-Ala-Ala and Pro-Pro-Pro in addition to the Gly-Gly-Ser linker (SE201-203). Each of these iterations maintained similar signaling potency and biochemical properties (**Figure S2b**), therefore Gly-Gly-Ser was chosen for the final linker sequence.

Finally, several mutations were introduced to the B-chain of single-chain relaxin-2 to improve its biochemical properties (SE204-206, **Figure 1b**). Met4 and Met25 were mutated to avoid residues prone to oxidation in the relaxin molecule. The Met residues were mutated to Lys according to the sequences of relaxin-2 orthologs, offering an idea of what residues may be tolerated in that position. Additionally, Trp28 was mutated to Ala in order to remove a residue that may increase protein polyreactivity. These three mutations were tolerated in the Fc–relaxin-2 fusions, having similar or better signaling potency than the native sequence (**Figure S2c**). In a docking model of the interactions between relaxin-2 and the ligand-binding ectodomain of the RXFP1 receptor (32), the residues targeted for mutagenesis are not positioned near the binding interface, explaining the lack of disruption to relaxin-2 activity (**Figure S3**).

### Biochemical and functional characterization of SE301

The optimized features of the Fc–relaxin-2 fusions were combined in the final molecule, SE301. This molecule contained, in order of sequence, the hemagglutinin signal sequence, the Fc N297Q fragment, a 3 residue Gly-Gly-Ser linker, and single-chain relaxin-2 with the redesigned “mini-C” linker and the Met4 to Lys, Met25 to Lys, and Trp28 to Ala mutations to the B-chain (**Figure 1b**). When tested in a Gs signaling assay, SE301 had an EC50 of 5.8 nM at human RXFP1, with an E_max_ at approximately 100% of native relaxin-2 (**Figure 1c**). While maintaining strong activation of RXFP1, SE301 also showed very little off-target activity at the related RXFP2 receptor (**Figure S4a**). The measured signaling potency of SE301 at human RXFP1 was confirmed with a secondary G_s_ signaling assay method, which showed close agreement with our initial experiments (EC_50_ of 7.1 nM, **Figure S5**). Next, the binding affinity for SE301 was tested using a flow cytometry assay with mammalian cells transfected with RXFP1 or empty vector. SE301’s K_D_ for human RXFP1 was determined to be 122 nM (**Figure 2a**), while a control molecule of the Fc N297Q fragment alone showed no binding (**Figure 2b**) or signaling (**Figure 1c**) with RXFP1-expressing cells.

**Figure 2:**
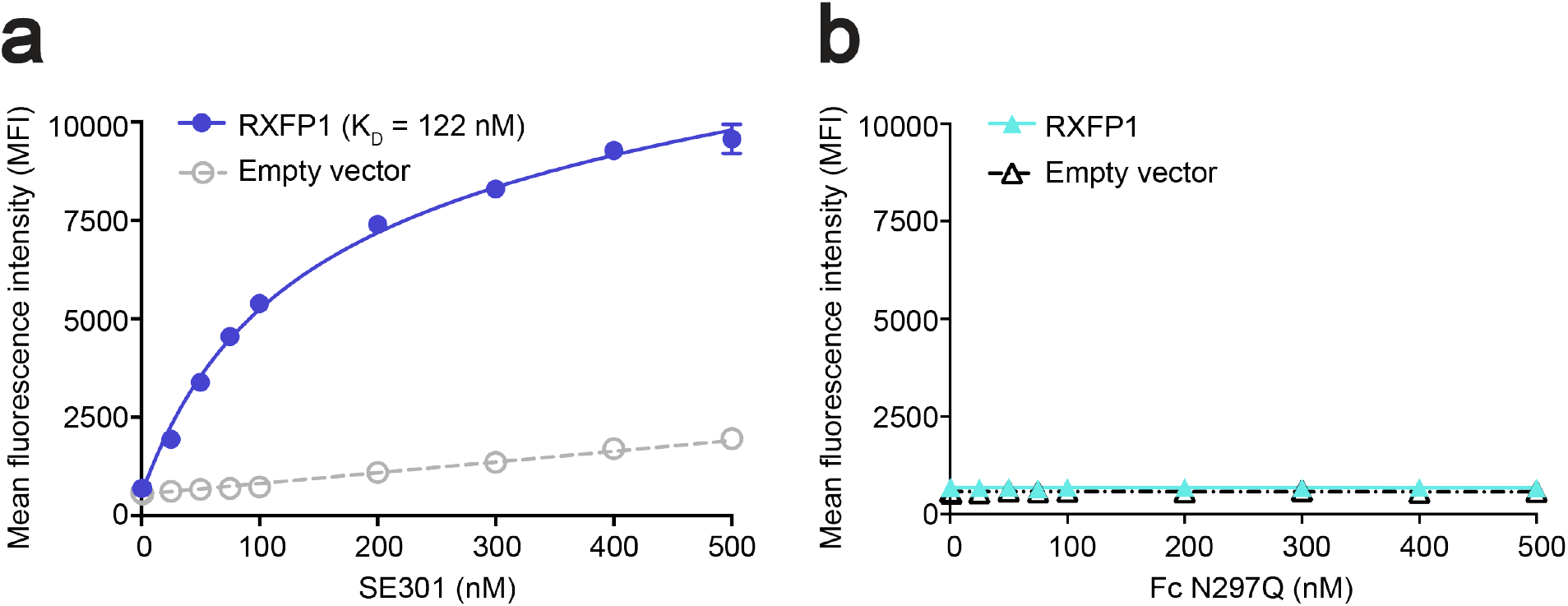
Determination of SE301 binding affinity for RXFP1. **a-b**, Flow cytometry binding data for SE301 (**a**) and Fc N297Q (**b**) using human RXFP1 and empty vector-transfected Expi293F cells. The K_D_ for SE301 at human RXFP1 was calculated to be 122 ± 36 nM. Data are mean ± s.e.m. from technical duplicates.

Additional biochemical studies for SE301 utilized differential scanning fluorimetry to determine a melting temperature (T_m_) of 57°C (**Figure 3d**). Furthermore, SE301 showed high stability at room temperature, maintaining similar signaling efficacy and potency after 4 weeks of incubation (**Figure 3b**). The protocols to produce SE301 as a secreted protein from mammalian cells utilized straightforward expression and purification methods, with a yield of approximately 150 mg per 1 L culture (See Methods). Collectively, the results from characterization studies showed that SE301 is a potent RXFP1 agonist with high purity, monodispersity, and yield (**Figure 3a,c**).

**Figure 3:**
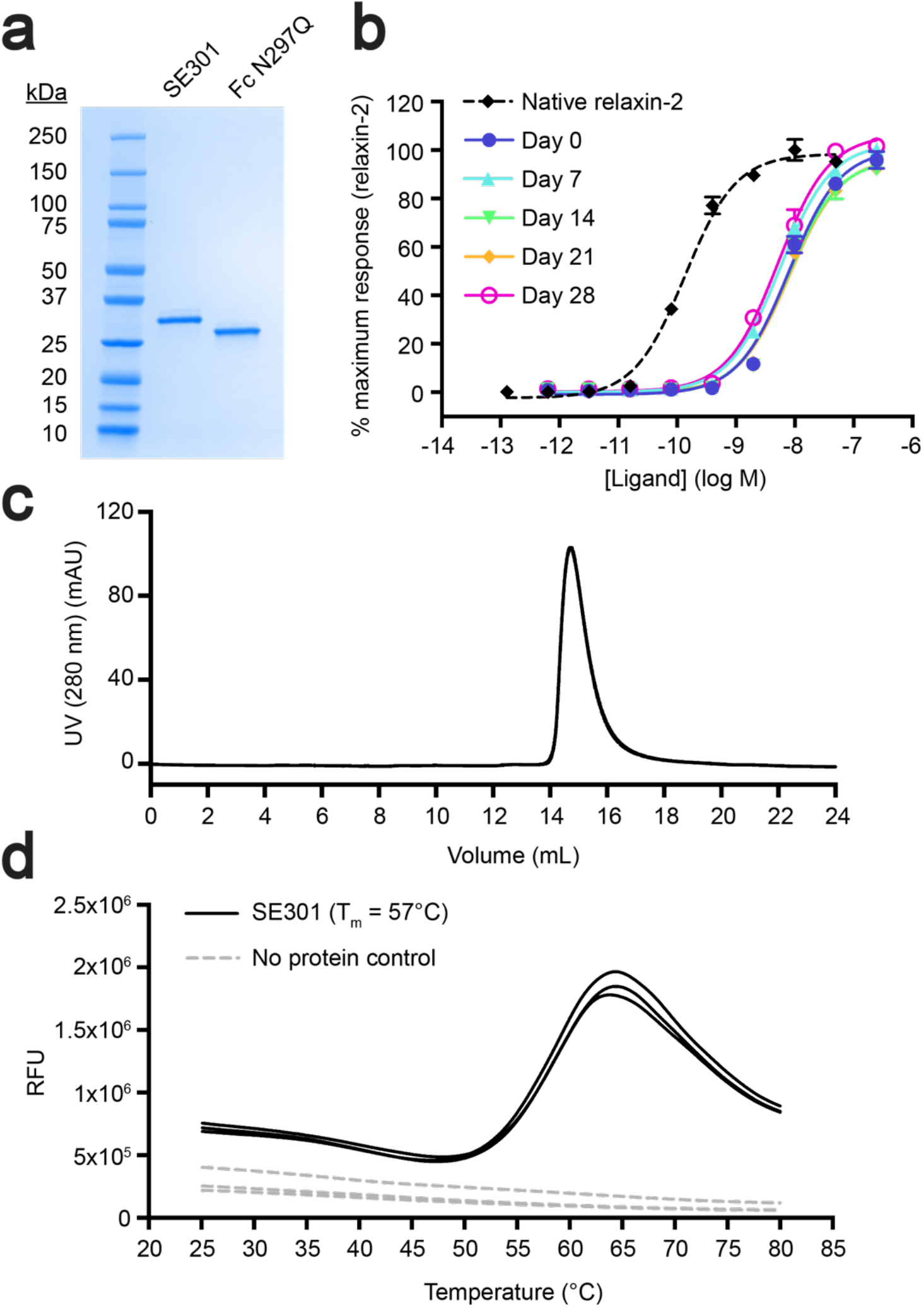
Characterization of SE301 purity, monodispersity, and stability. **a**, Coomassie-stained SDS-PAGE gel of SE301 and Fc N297Q. **b**, CRE-SEAP G_s_ signaling testing the activity of SE301 at human RXFP1 after incubations at room temperature for 0, 7, 14, 21, and 28 days, showing that SE301 retains activity after 4 weeks at room temperature. Data are normalized to the native relaxin-2 response at human RXFP1 and are mean ± s.e.m. from technical triplicates. **c**, Size exclusion profile for SE301 shows a monodisperse peak. **d,** Differential scanning fluorimetry for SE301 determined its melting temperature to be 57°C.

### Pharmacokinetics of SE301

After biochemical and functional characterization of the SE301 molecule, we conducted experiments to determine the serum half-life of SE301 *in vivo*. To answer this question, we conducted a pharmacokinetics study in mice using a single injection of SE301 at one of three doses, 1, 5, or 50 mg/kg. Serum samples were taken before injection, and then at 2, 24, 72, and 168 hours post-injection. To determine the concentration of SE301 remaining in circulation at each timepoint, the serum samples were analyzed by an ELISA detecting the human IgG1 Fc of SE301. Based on concentrations interpolated from the ELISA data, the serum half-life was calculated to be between 77.5 and 130 hours, depending on the dose of SE301 (**Figure 4a**). These results approach the 6 to 8-day serum half-life of IgGs in mice (33), showing that the Fc fragment was able to confer a longer circulating half-life to SE301.

**Figure 4:**
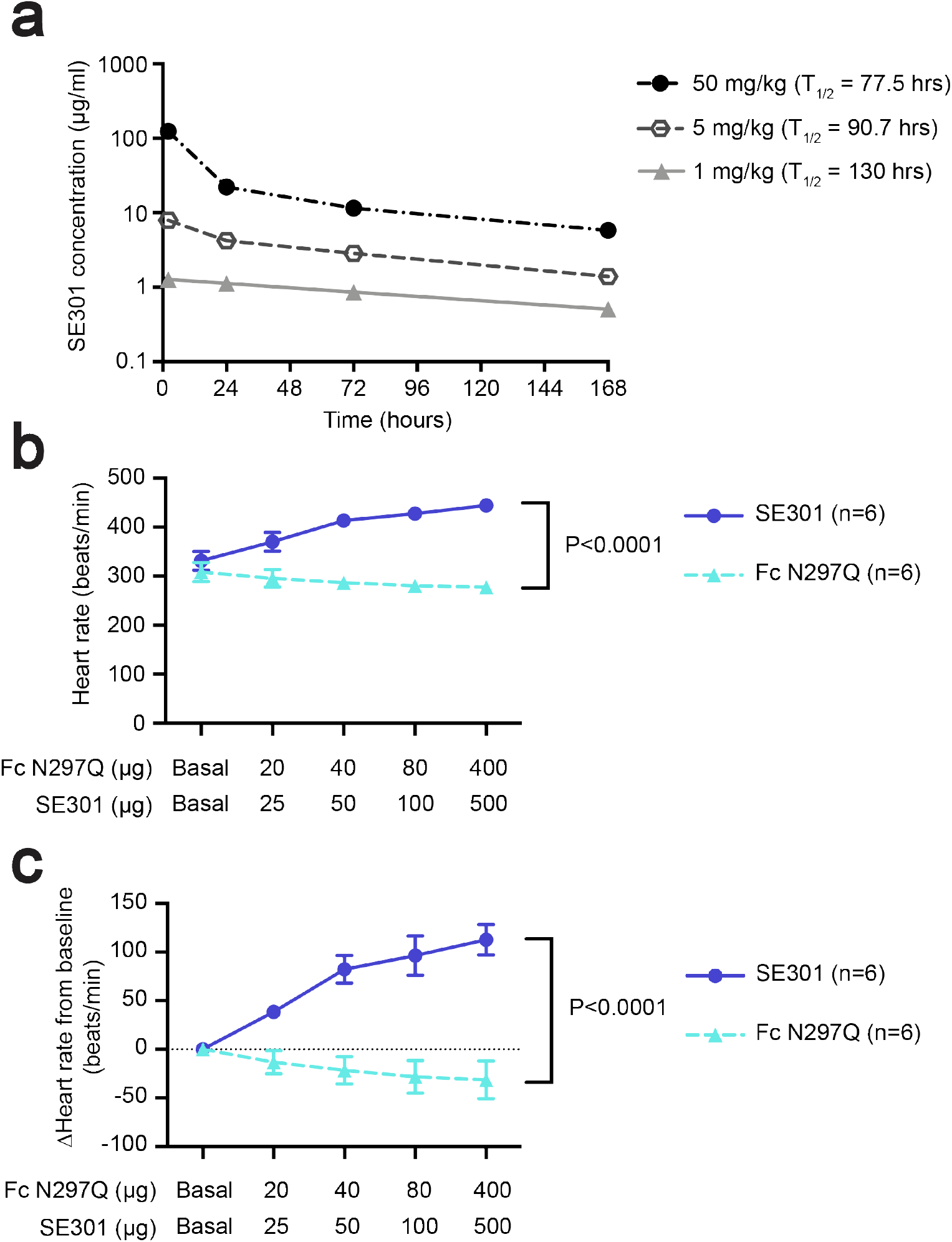
Pharmacokinetics and *in vivo* hemodynamics. **a**, Pharmacokinetics study of SE301 in mice used a single intraperitoneal injection and measured the SE301 serum concentration by ELISA from samples taken before injection, and 2, 24, 72, and 168 hours post-injection. The serum half-life for SE301 was calculated to be between 77.5 to 130 hours. Data are mean ± s.e.m. and n=3 for each dose. **b-c**, Mouse hemodynamics study for SE301 or Fc N297Q injection in mice. Heart rate was monitored upon increasing doses of SE301 or Fc N297Q. Plots show either the measured heart rate (**b**) or calculated as a change from the baseline heart rate (**c**). Data are mean ± s.e.m. and n=6, p<0.0001.

### SE301 shows *in vivo* activity

After establishing the long serum half-life of SE301, we conducted a study to determine the activity of the molecule at the RXFP1 receptor *in vivo*. In rodents, RXFP1 expressed in the atria of the heart causes a positive chronotropic effect upon relaxin-2 treatment, with an increase in heart rate of around 130 beats per minute (bpm) above baseline (34,35). The heart rate changes in response to relaxin-2 directly result from activation of RXFP1 and are not an effect of catecholamine release (35). The chronotropic effects do not translate to humans because RXFP1 is not expressed in the atria, and relaxin-2 administration has been shown to have no effect on heart rate in clinical trials (36). Although it does not directly translate to any human disease, the effect on rodent heart rate represents a short-term experiment testing RXFP1 agonist activity *in vivo*. Therefore, we chose to carry out a mouse hemodynamics study to test SE301, in advance of longer-timeline studies with animal models of human disease.

First, the ability of our engineered human relaxin-2 fusion to activate mouse RXFP1 was tested using a cell-based G_s_ signaling. The assay determined an EC_50_ of 8.6 nM for SE301 at mouse RXFP1, establishing the utility of mouse models for our *in vivo* studies (**Figure S4b**). In the hemodynamics study, mice were anesthetized and the heart rate of the left ventricle was monitored. Increasing doses of the Fc N297Q alone control molecule or SE301 were administered by intravenous injection at 10-minute intervals. Mice showed a dose-dependent increase in heart rate in response to SE301, increasing from around 330 bpm at baseline to 445 bpm after injection with 500 μg of SE301 (**Figure 4b,c**). In contrast, the mice showed no increase in heart rate in response to Fc N297Q. Together, these data established the ability of SE301 to activate the RXFP1 receptor *in vivo*.

## Discussion

Here we described each step in the design of a long half-life RXFP1 agonist. Through applying protein engineering strategies to the hormone relaxin-2, we generated a potent agonist of the RXFP1 receptor with a long serum half-life, as well as high yield, purity, and biochemical stability. Our general approach should be extensible to other relaxin family peptide hormones; although optimal mutations, linker lengths, and fusion sites will likely vary due to the different modes of ligand recognition among the relaxin receptors.

Mouse studies with our final Fc–relaxin-2 fusion, SE301 confirmed an extended half-life of between 3 to 5 days and established its *in vivo* activity at RXFP1. In a mouse hemodynamics study, SE301 increased heart rate in a manner comparable with the native relaxin-2 peptide (35). Future work will test the efficacy of SE301 in animal models of cardiovascular or fibrotic conditions in which treatment with the native relaxin-2 peptide has proven promising. In those experiments, recombinant versions of native relaxin-2 are typically administered continuously (9–11,13), similar to the intravenous infusions of the relaxin-2 peptide used in clinical trials (14). Given its extended half-life, our engineered Fc–relaxin-2 has the potential to achieve a similar improvement in disease phenotypes through weekly or biweekly administration by subcutaneous injections. The results of these experiments will potentially provide a path toward accessing the beneficial biological effects of relaxin-2 for a wider range of indications.

## Acknowledgements

We would like to thank Dr. Kelly L. Arnett for assistance with differential scanning fluorimetry measurements and the Center for Macromolecular Interactions at Harvard Medical School where differential scanning fluorimetry experiments were performed. This work was supported by National Institutes of Health grants HL056687 and HL075443 to H.A.R., an NIH Ruth L. Kirschstein predoctoral fellowship (F31 GM128233) to S.C.E., and a Blavatnik Biomedical Accelerator grant from Harvard Medical School to A.C.K.

## Competing interests statement

A.C.K. and S.C.E are inventors on a patent application for engineered single-chain relaxin proteins. A.C.K. is a co-founder and consultant for Tectonic Therapeutic and Seismic Therapeutic and for the Institute for Protein Innovation, a non-profit research institute.

## Methods

### Molecular cloning

DNA encoding single-chain relaxins with an N-terminal hemagglutinin signal sequence and C-terminal 6x His-tag were cloned into the pcDNA-Zeo-tetO vector (37) using PCR and NEBuilder HiFi DNA Assembly Mix (New England Biolabs). Fc fusion single-chain relaxins and the Fc N297Q negative control were cloned into pcDNA-Zeo-tetO with an N-terminal hemagglutinin signal sequence. For human and mouse RXFP1 and human RXFP2 expression constructs, receptors were cloned into pcDNA-Zeo-tetO with an N-terminal hemagglutinin signal sequence and FLAG tag.

### Protein expression and purification

#### His-tagged proteins

His-tagged relaxins were expressed as secreted proteins in Expi293F cells containing a stably integrated tetracycline repressor (Expi293F tetR, Thermo Fisher Scientific) grown in Expi293 media (Thermo Fisher Scientific). Cells were transiently transfected with polyethylenimine, enhanced 24 hours post-transfection with 0.4% glucose, 5 mM sodium butyrate, and 3 mM sodium valproic acid, and induced 48 hours post-transfection with 5 mM sodium butyrate and 4 μg/mL doxycycline. Supernatant containing the single-chain relaxins was harvested from the cultures 5 days after induction by centrifugation at 4,000 ×g for 15 minutes at 4°C.

To purify His-tagged single-chain relaxins, supernatant was filtered with a glass fiber filter and loaded over Nickel Excel resin (GE Healthcare) equilibrated with 30 mM MES pH 6.5, 300 mM sodium chloride. The resin was washed with 30 mM MES pH 6.5, 300 mM sodium chloride, 20 mM imidazole, and protein was eluted with 30 mM MES pH 6.5, 300 mM sodium chloride, 500 mM imidazole. Ammonium sulfate was added to the eluted protein to 60% saturation and rotated at 4°C for 1 hour. Precipitated protein was centrifuged at 10,000 ×g, for 15 minutes at 4°C and the pellet was resuspended in 30 mM MES pH 6.5, 300 mM sodium chloride. Resuspended protein was filtered using a 0.1 um pore size centrifugal filter and loaded onto a Sephadex S200 column (GE Healthcare) for size exclusion chromatography (SEC). Peak fractions were collected and concentrated with a centrifugal concentrator with a 3 kDa molecular weight cut off. Purity of the proteins was assessed by SDS-PAGE gel. Aliquots were flash frozen in liquid nitrogen and stored at −80°C.

#### Fc fusion proteins

Fc fusion single-chain relaxins and the Fc N297Q control were expressed in Expi293F tetR cells as stated above. Supernatant containing the Fc fusions was harvested from the cultures 5 days after induction by centrifugation at 4,000 ×g for 15 minutes at 4°C. Supernatant containing the Fc fusions was diluted 1:1 in 20 mM HEPES pH 7.5, 150 mM sodium chloride (HBS) and loaded onto protein G resin (GE Healthcare) equilibrated with HBS. The resin was washed with HBS and protein was eluted with 100 mM glycine pH 2.5. The elution was neutralized to pH 7.5 with HEPES and dialyzed overnight in HBS. Samples prepared for mouse studies were dialyzed into phosphate buffered saline (PBS) at 4°C. Elutions from large-scale cultures were diluted in 100 mM glycine pH 2.5 prior to neutralization to avoid precipitation upon pH change. Dialyzed protein was concentrated with a centrifugal concentrator with a 3 kDa molecular weight cut off. SDS-PAGE gels and analytical SEC were used to analyze proteins for purity and monodispersity. Proteins were aliquoted, flash frozen in liquid nitrogen, and stored at −80°C.

### Cellular assays

#### CRE-SEAP

G_s_ signaling was measured using an assay that indirectly detects cAMP production through transcription of the reporter enzyme secreted embryonic alkaline phosphatase (SEAP) (38). Briefly, clear 96-well plates were coated with 30 uL of 10 ug/mL poly-D-lysine, washed with PBS, and HEK293T cells (ATCC) were plated at 2.4×10^4^ cells/well in Dulbecco’s Modified Eagle Medium (DMEM) with 10% (v/v) fetal bovine serum (FBS). The next day, the medium was replaced with 50 uL of serum-free DMEM. Lipofectamine 2000 (Thermo Fisher Scientific) was used to transfect cells at 70% confluency with 20 ng of CRE-SEAP reporter plasmid (Clontech) and 20 ng of receptor or empty vector pcDNA-Zeo-tetO DNA per well. Transfections were incubated for 5 hours at 37°C, then the medium was replaced with 200 uL of serum-free DMEM plus ligand dilution curves. Twenty-four hours later, the plates were incubated at 70°C for 2 hours. A solution of the SEAP substrate, 4-methylumbelliferyl phosphate (Sigma Aldrich), was prepared at 120 uM in 2M diethanolamine bicarbonate pH 10. The substrate solution was mixed with an equal volume (100 uL) of supernatant and incubated at room temperature for 15 minutes. An Envision 2103 Multilabel Reader (Perkin Elmer) was used to measure fluorescence with an excitation wavelength of 360 nm and an emission wavelength of 449 nm. Signaling was calculated as a percentage of native relaxin-2 response on either human or mouse RXPF1 and plotted using GraphPad Prism.

#### GloSensor

A real-time, live-cell signaling assay was used as a second method to measure SE301 activation of Gs signaling through RXFP1. The assay was carried out as previously described (32). Briefly, white, clear-bottom 96-well plates were coated with poly-D-lysine and washed with PBS. HEK293T cells were then plated at 2.0×10^4^ cells/well. The next day, cells were transfected with human RXFP1 pcDNA-Zeo-tetO and the GloSensor reporter plasmid using FuGENE (Promega), according to the manufacturer’s instructions. The cells were incubated for 24 hours at 37°C with 5% CO_2_. The next day, the media was changed to 40 μL CO_2_-independent media (Thermo Fisher Scientific) with 10% (v/v) FBS and 2 mg/mL D-luciferin (Goldbio). The plates were then incubated for 2 hours at room temperature (RT) in the dark. After 2 hours, luminescence was measured before adding ligands using a SpectraMax M5 microplate reader with a 1 second integration time. A dilution series of SE301 or native relaxin-2 were added to the cells and the luminescence measurement was repeated at 5, 10, 15, 20, 25, and 30 minutes after ligand addition. Signaling was calculated as a percentage of native relaxin-2 response on human RXPF1 and plotted using GraphPad Prism.

### Differential scanning fluorimetry

For differential scanning fluorimetry, SE301 was dialyzed overnight into PBS. Samples were prepared with 0.1 mg/mL SE301 or dialysis buffer control and mixed with Protein Thermal Shift Dye (Applied Biosystems) in a 1:100 ratio (v/v) of protein to dye in 96-well plates (Applied Biosystems). Differential scanning fluorimetry was carried out using the Life Technologies Quant Studio 6 with temperatures from 25 to 99°C, increasing by 3°C per minute. Fluorescence was detected with 470 nm excitation and 586 nm emission filters. The fluorescence readings as a function of temperature were analyzed in the Protein Thermal Shift Software (Applied Biosystems) using the Boltzmann equation and plotted using GraphPad Prism.

### Flow cytometry binding assay

To measure SE301 binding affinity, flow cytometry was used with Expi293F cells transfected with human RXFP1 or empty pcDNA-Zeo-tetO vector. Expi293F tetR cells were grown in Expi293 media and transfected using FectoPRO (Polyplus), according to the manufacturer’s protocols. The cells were enhanced 24 hours post-transfection with 0.4% glucose and induced 48 hours post-transfection with 4 μg/mL doxycycline and 5 mM sodium butyrate. After 24 hours of induction, cells were harvested by spinning at 200 ×g for 5 minutes at 4°C and washed once with HBS with 1% (v/v) FBS and 2 mM calcium chloride (Binding Buffer). Cells were plated into a V-bottom 96-well plate (Corning) at 100,000 cells/well and blocked by incubation in Binding Buffer for 30 minutes at 4°C. After blocking, cells were centrifuged at 200 ×g for 5 minutes at 4°C, resuspended in 100 μL of Binding Buffer containing a dilution series of SE301 or Fc N297Q, and incubated for 1 hour at 4°C. Cells were then centrifuged at 200 xg for 5 minutes at 4°C, washed twice with 200 μL Binding Buffer, and resuspended in 100 μL Binding Buffer containing 100 nM M1 anti-FLAG antibody labeled with Alexa Fluor 488 and Alexa Fluor 647 anti-human IgG Fc (BioLegend) diluted 1:100 (v/v). Cells were incubated in secondary antibodies for 30 minutes at 4°C, washed once with 200 μL Binding Buffer, and resuspended in 100 μL Binding Buffer for flow cytometry. Samples were analyzed on a BD Accuri C6 flow cytometer (BD Biosciences) and gated according to plots of FSC-A/SCA-A, FSC-A/FSC-H, and receptor expression according to Alexa Fluor 488 M1 anti-FLAG antibody binding. Approximately 1000 events/sample were collected from cells expressing receptor for human RXFP1-transfected cells or post-FSC-A/FSC-H gating for empty vector-transfected cells. Mean fluorescence intensities for Alexa Fluor 647 anti-human IgG Fc binding were plotted and analyzed in GraphPad Prism.

### Mouse pharmacokinetics study

A pharmacokinetics study in mice was conducted to determine the serum half-life of the SE301 molecule. SE301 was prepared in sterile PBS at 10 mg/mL, using methods described above. For the study, male CD-1 mice were used with intraperitoneal injections (IP) of SE301 at one of three doses, 1, 5, or 50 mg/kg. Nine mice were used in the pharmacokinetic study, three per each dose of SE301, and each was between 7 to 10 weeks of age and weighed between 29 and 40 grams. The IP injection of SE301 was administered via hypogastric regions, and blood samples were taken from the mice before dosing, and 2, 24, 72, and 168 hours post-dosing. At each timepoint, at least 0.6 mL of blood was collected from each animal. The blood samples were stored at room temperature for about 30 minutes, and then were centrifuged at 2,500 ×g for 15 minutes at 4°C. After centrifugation, the serum was collected, an aliquot was taken for analysis, and the samples were frozen over dry ice and stored at −60°C or lower. An ELISA was conducted detecting human IgG1 Fc to determine the amount of SE301 per sample. Serum SE301 concentration versus timepoint data were plotted in the WinNonlin software program to derive pharmacokinetic parameters.

### Mouse hemodynamics study

Animal experiments carried out for this study were handled according to approved protocols and animal welfare regulations mandated by the Institutional Animal Care and Use Committee of Duke University Medical Center. Eight to twelve-week-old C57BL/6J wild-type mice of both sexes were used for this study. Mice were anesthetized with ketamine (100 mg/kg) and xylazine (2.5 mg/kg), and bilateral vagotomy was performed. The left ventricle blood pressure and heart rate were monitored with a 1.4 French (0.46 mm) high fidelity micromanometer catheter (ADInstruments) connected to a pressure transducer (ADInstruments). Basal blood pressure was recorded at steady state after catheter insertion (2-3 min after insertion). Graded doses of Fc N297Q (20, 40, 80, 400 μg) or SE301 (25, 50, 100, 500 μg) were administered at 10 min intervals by intravenous injection through a jugular vein. The blood pressure was monitored continuously and recorded at the steady state (10 min after each injection). Data analysis was performed using LabChart 8 software (ADInstruments).

