## Supplementary_information for "Engineering and characterization of a long half-life relaxin receptor RXFP1 agonist"

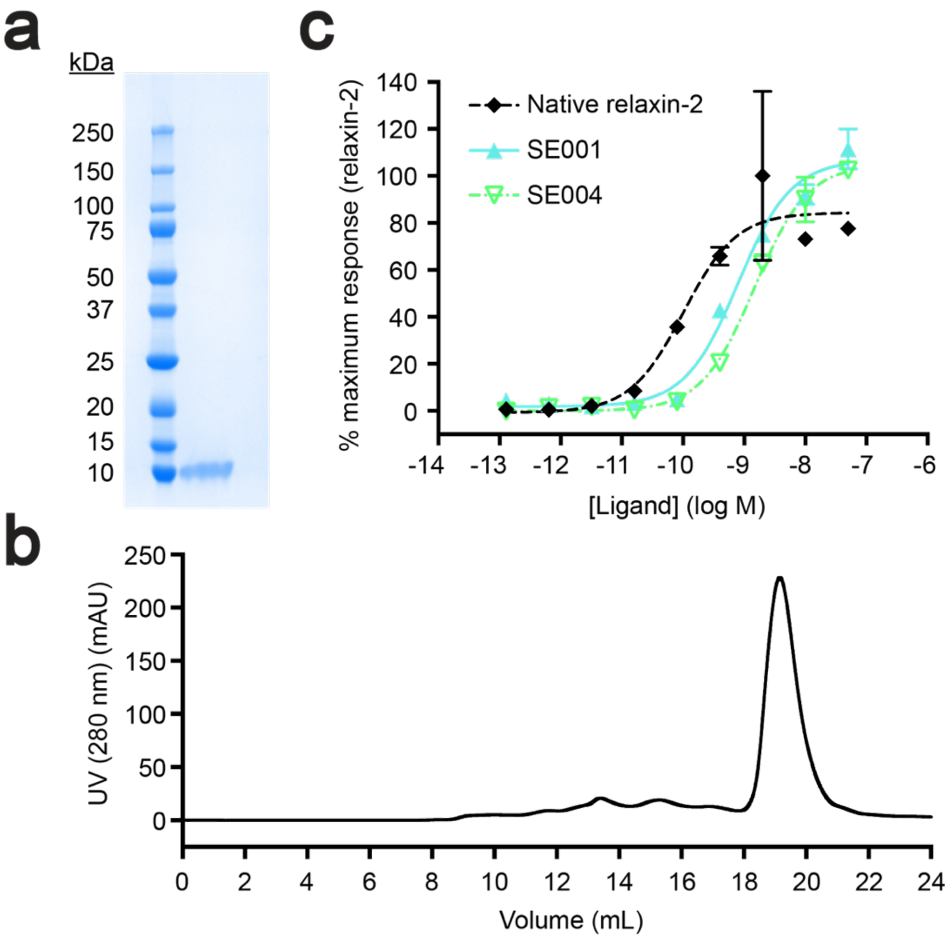


**Figure S1: Purification and characterization of single-chain relaxin-2. a**, Coomassie-stained SDS-PAGE gel for His-tagged single-chain relaxin-2 (SE001). **b**, Size exclusion chromatography profile for His-tagged single-chain relaxin-2 (SE001). **c**, CRE-SEAP $G_s$ signaling assay with human RXFP1 for His-tagged single-chain relaxin-2 (SE001) and His-tagged, protein C-tagged single-chain relaxin-2 (SE004). Data are normalized to the native relaxin-2 response and are mean ± s.e.m. from technical triplicates.

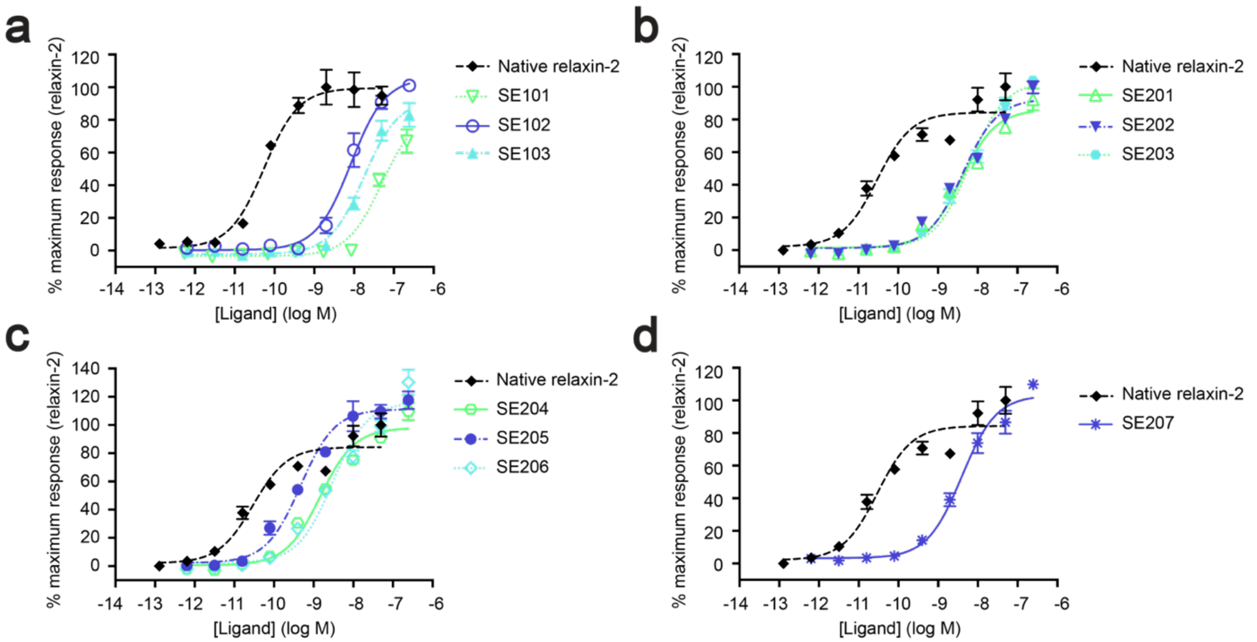


**Figure S2: Optimizations of Fc–relaxin-2 fusions. a-d**, CRE-SEAP $G_s$ signaling data for human RXFP1 using native relaxin-2 compared to SE101–SE103 (**a**), SE201–SE203 (**b**), SE204–SE206 (**c**), and SE207 (**d**). Data are normalized to the native relaxin-2 response at human RXFP1 and are mean ± s.e.m. from technical triplicates.

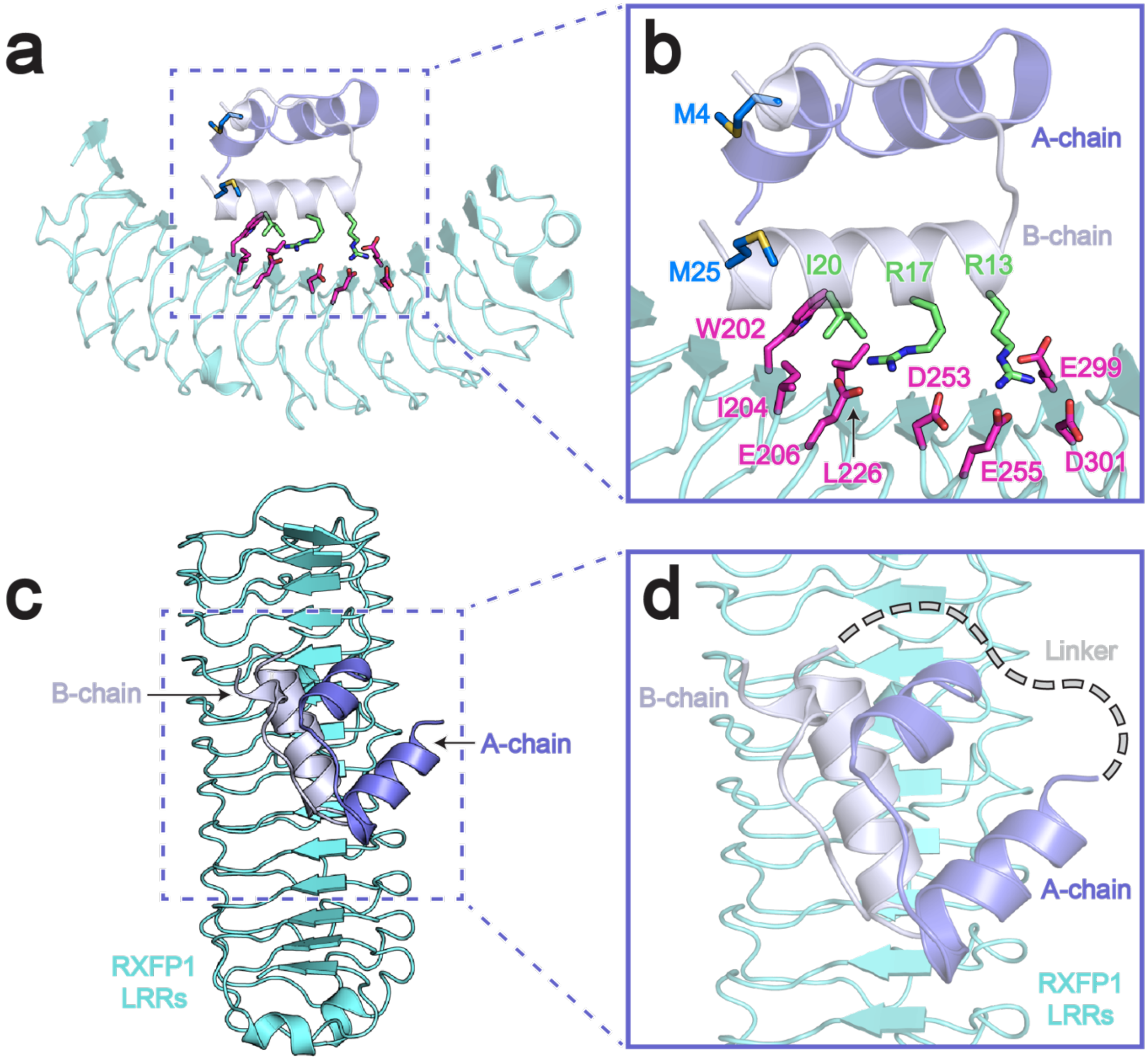


**Figure S3: Sites of relaxin-2 engineering in Fc–relaxin-2 fusion constructs. a**,**c**, Docking model of relaxin-2 bound to the leucine-rich repeats (LRRs) of RXFP1’s ectodomain (1). **b**, Details of the relaxin-2–LRR interface. In magenta are RXFP1 residues involved in the binding interface, in green are the “relaxin-binding cassette” residues of relaxin-2’s B-chain, Arg13, Arg17, and Ile20, and in blue are the Met residues mutated to Lys in SE301. The Trp28 residue was not included in the docking model. The model shows that the Met residues on the relaxin-2 B-chain are not positioned near the binding interface. **d**, Based on the model, the position of the single-chain relaxin-2 “mini-C” linker likely does not interfere with the binding of relaxin-2 to RXFP1.

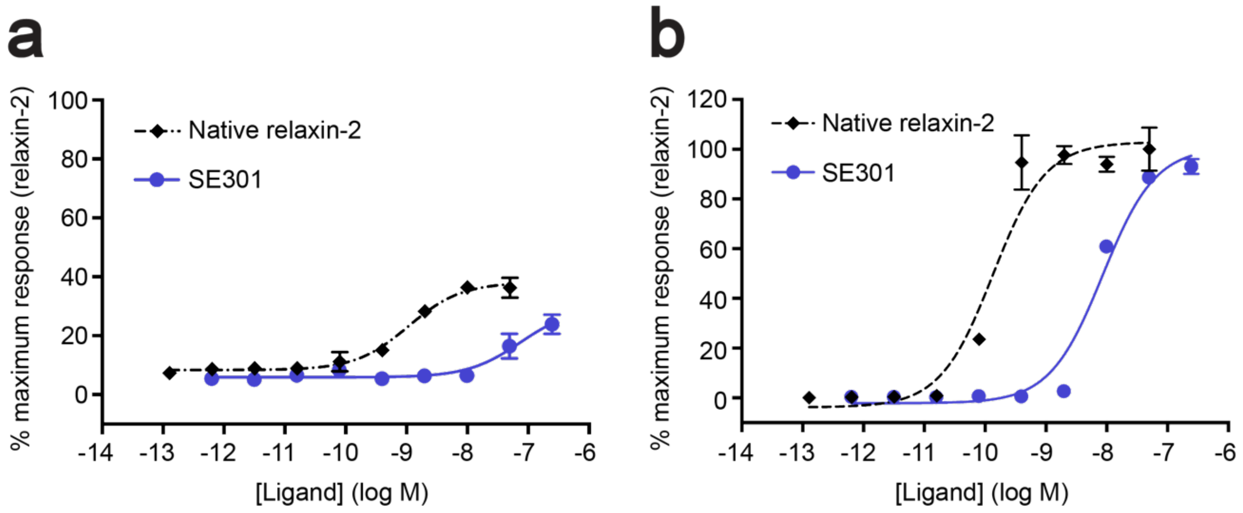


**Figure S4: SE301 signaling activity at human RXFP2 and mouse RXFP1. a**, CRE-SEAP $G_s$ signaling assay data for SE301 compared to native relaxin-2 at human RXFP2. Data are normalized to the native relaxin-2 response at human RXFP1 and are mean ± s.e.m. from technical triplicates. **b**, CRE-SEAP $G_s$ signaling assay data for SE301 compared to native human relaxin-2 at mouse RXFP1. Data are normalized to the native human relaxin-2 response at mouse RXFP1 and are mean ± s.e.m. from technical triplicates.

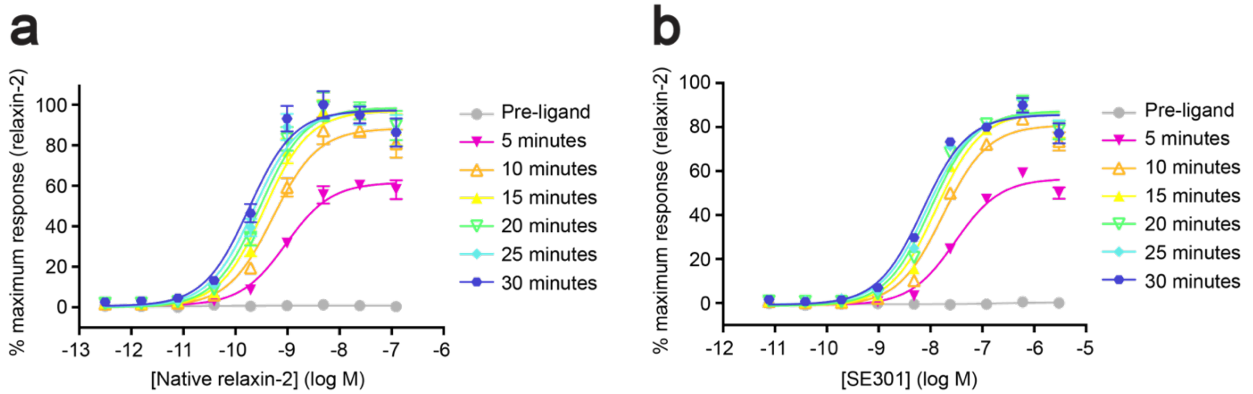


**Figure S5: SE301 versus native relaxin-2 activity in the $G_s$ GloSensor cAMP assay. a-b**, GloSensor $G_s$ signaling assay data for native relaxin-2 (**a**) and SE301 (**b**) with measurements taken before ligand addition, and 5, 10, 15, 20, 25, and 30 minutes after ligand addition. Data are normalized to the native relaxin-2 response at human RXFP1 and are mean ± s.e.m. from technical triplicates.

**Table S1: CRE-SEAP $G_s$ signaling assay $EC_{50}$ and $E_{max}$ data for Figure 1c**

†Mean ± s.e.m., n=3 technical replicates.

| Ligand | $pEC_{50}$ | $E_{max}$ (%) |
|---|---|---|
| Native relaxin-2 | 10.0 ± 0.06 | 100 ± 2.0 |
| SE301 | 8.2 ± 0.05 | 110 ± 2.5 |
| Fc N297Q | ND | ND |

**Table S2: CRE-SEAP $G_s$ signaling assay $EC_{50}$ and $E_{max}$ data for Figure S1c**

†Mean ± s.e.m., n=3 technical replicates.

| Ligand | $pEC_{50}$ | $E_{max}$ (%) |
|---|---|---|
| Native relaxin-2 | 10.0 ± 0.2 | 100 ± 6.9 |
| SE001 | 9.1 ± 0.1 | 126 ± 3.3 |
| SE004 | 8.8 ± 0.1 | 124 ± 2.9 |

**Table S3: CRE-SEAP $G_s$ signaling assay $EC_{50}$ and $E_{max}$ data for Figure S2**

†Mean ± s.e.m., n=3 technical replicates.

| Ligand | $pEC_{50}$ | $E_{max}$ (%) |
|---|---|---|
| Native relaxin-2 | 10.3 ± 0.1 | 100 ± 3.1 |
| SE101 | 7.3 ± 0.1 | 86 ± 7.4 |
| SE102 | 8.1 ± 0.1 | 106 ± 3.9 |
| SE103 | 7.7 ± 0.1 | 93 ± 4.5 |
| Native relaxin-2 | 10.5 ± 0.1 | 100 ± 3.5 |
| SE201 | 8.4 ± 0.1 | 102 ± 3.1 |
| SE202 | 8.4 ± 0.1 | 110 ± 3.3 |
| SE203 | 8.2 ± 0.1 | 122 ± 3.0 |
| SE204 | 8.8 ± 0.1 | 116 ± 3.3 |
| SE205 | 9.3 ± 0.1 | 132 ± 3.1 |
| SE206 | 8.5 ± 0.1 | 138 ± 4.9 |
| SE207 | 8.4 ± 0.1 | 122 ± 3.3 |

**Table S4: CRE-SEAP $G_s$ signaling assay $EC_{50}$ and $E_{max}$ data for Figure S4**

[†]Mean ± s.e.m., n=3 technical replicates.

| Ligand | $pEC_{50}$ | $E_{max}$ (%) |
|---|---|---|
| Native relaxin-2, human RXFP2 | 9.0 ± 0.1 | 38 ± 1.7 |
| SE301, human RXFP2 | 7.1 ± 0.3 | 30 ± 5.7 |
| Native relaxin-2, mouse RXFP1 | 9.9 ± 0.1 | 100 ± 4.2 |
| SE301, mouse RXFP1 | 8.1 ± 0.1 | 98 ± 3.7 |

**Table S5: GloSensor $G_s$ signaling assay $EC_{50}$ and $E_{max}$ data for Figure S5**

[†]Mean ± s.e.m., n=3 technical replicates.

| Ligand | $pEC_{50}$ | $E_{max}$ (%) |
|---|---|---|
| Native relaxin-2, Pre-ligand | ND | 1 ± 0.2 |
| Native relaxin-2, 5 minutes | 9.0 ± 0.1 | 63 ± 1.7 |
| Native relaxin-2, 10 minutes | 9.3 ± 0.1 | 91 ± 2.7 |
| Native relaxin-2, 15 minutes | 9.4 ± 0.1 | 100 ± 2.9 |
| Native relaxin-2, 20 minutes | 9.5 ± 0.1 | 101 ± 2.9 |
| Native relaxin-2, 25 minutes | 9.6 ± 0.1 | 100 ± 3.2 |
| Native relaxin-2, 30 minutes | 9.7 ± 0.1 | 100 ± 3.1 |
| SE301, Pre-ligand | ND | 0 ± 0.5 |
| SE301, 5 minutes | 7.5 ± 0.1 | 58 ± 1.6 |
| SE301, 10 minutes | 7.8 ± 0.1 | 83 ± 2.0 |
| SE301, 15 minutes | 7.9 ± 0.1 | 89 ± 2.0 |
| SE301, 20 minutes | 8.0 ± 0.1 | 90 ± 2.2 |
| SE301, 25 minutes | 8.1 ± 0.1 | 89 ± 2.2 |
| SE301, 30 minutes | 8.1 ± 0.1 | 88 ± 2.0 |

**Table S6: CRE-SEAP $G_s$ signaling assay $EC_{50}$ and $E_{max}$ data for Figure 3b**

†Mean ± s.e.m., n=3 technical replicates.

| **Ligand** | **$pEC_{50}$** | **$E_{max}$ (%)** |
|---|---|---|
| Native relaxin-2 | 9.9 ± 0.05 | 100 ± 1.7 |
| SE301 Day 0 | 8.1 ± 0.05 | 102 ± 2.4 |
| SE301 Day 7 | 8.2 ± 0.03 | 104 ± 1.3 |
| SE301 Day 14 | 8.1 ± 0.1 | 98 ± 2.4 |
| SE301 Day 21 | 8.1 ± 0.04 | 98 ± 2.0 |
| SE301 Day 28 | 8.3 ± 0.04 | 108 ± 2.1 |
